# Mapping lung hematopoietic progenitors: Developmental kinetics and response to Influenza A viral infection

**DOI:** 10.1101/2023.10.09.561586

**Authors:** Kyle T. Mincham, Jean-Francois Lauzon-Joset, James F. Read, Patrick G. Holt, Philip A. Stumbles, Deborah H. Strickland

**Affiliations:** Wal-yan Respiratory Research Centre, Telethon Kids Institute, Nedlands, Western Australia, Australia; National Heart and Lung Institute, Imperial College London, London, United Kingdom; Centre de recherche de l’Institut de Cardiologie et de Pneumologie de Québec, Université Laval, Québec, Canada; Asthma and Airway Disease Research Center, University of Arizona, Tucson, Arizona, United States; Child Health Research Centre, The University of Queensland, Brisbane, Queensland, Australia; Medical, Molecular and Forensic Sciences, Murdoch University, Perth, Western Australia, Australia

**Author notes:** **Funding:** This study was funded by the National Health and Medical Research Council of Australia.

**Keywords:** Hematopoietic progenitors, hematopoiesis, granulocyte-macrophage, extravascular lung, bone marrow, Influenza, fetal, extramedullary

## Abstract

The bone marrow is a specialised niche responsible for the maintenance of hematopoietic stem and progenitor cells during homeostasis and inflammation. Recent studies however have extended this essential role to the extramedullary and extravascular lung microenvironment. Here, we provide further evidence for a reservoir of hematopoietic stem and progenitor cells within the lung from embryonic day 18.5 until adulthood. These lung progenitors display distinct microenvironment-specific developmental kinetics compared to their bone marrow counterparts, exemplified by a rapid shift from a common myeloid to megakaryocyte-erythrocyte progenitor dominated niche with increasing age. In adult mice, Influenza A viral infection results in a transient reduction in multipotent progenitors within the lungs, with a parallel increase in downstream granulocyte-macrophage progenitors and dendritic cell populations associated with acute viral infections. Our findings suggest lung hematopoietic progenitors play a role in re-establishing immunological homeostasis in the respiratory mucosa, which may have significant clinical implications for maintaining pulmonary health following inflammatory perturbation.

## Introduction

Hematopoiesis is an evolutionarily conserved process, with primitive hematopoietic progenitors first appearing in the yolk sac during early embryogenesis (1, 2). This primal development is rapidly proceeded by a sequential wave of definitive hematopoietic stem and progenitor cells (HSPC), which migrate to colonise the fetal liver, spleen and eventually the bone marrow (BM) (3). Once established, the BM is considered the dominant niche responsible for postnatal maintenance of HSPCs.

The classic hierarchical model of HSPC commitment within the BM involves the progression of lineage negative (Lin^-^) Sca-1^+^c-Kit^+^ (termed LSK cells) hematopoietic stem cells (HSC) through multiple intermediate stages of development, which ultimately lose their self-renewal capacity following upregulation of fms-like tyrosine kinase 3 (Flt3) expression (4, 5). In response to microenvironmental cues, multipotent progenitors (MPP) commit to oligopotent common lymphoid progenitors (CLP) (6), giving rise to T cell, B cell, innate lymphoid cell progenitor and pre-natural killer progenitor (pre-NKp) subsets (7). Simultaneously, all myeloid progenitors (MP), which arise from common myeloid progenitors (CMP), commit to megakaryocyte-erythrocyte progenitors (MEP), granulocyte macrophage progenitors (GMP), and their downstream macrophage-dendritic cell progenitors (MDP) (8, 9). In addition to the BM, similar niches containing tissue-specific stem cells with capacity for long-term self-renewal have been identified in multiple peripheral tissues, including the spleen (10), lymph nodes (11), adipose tissue (12, 13) and lung (14-16), shedding light on extramedullary hematopoiesis in homeostatic and pathogenic regulation.

Throughout development, the fetal hematopoietic and pulmonary systems undergo simultaneous organogenesis (17). Convergence of these two processes occurs as early as embryonic (E) day 16.5 in mice, with fetal liver-derived monocytes seeding the lung to establish a self-renewing population of alveolar macrophages, which is maintained throughout life independent of the BM (18, 19). Morphogenic and immunological maturation of the airways, however, is not achieved until many years later. As such, early progenitors likely dictate critical functions in neonatal protection, while disruption of their developmental kinetics may have consequential effects on future health (20).

In this regard, maintenance of immunological homeostasis in the respiratory mucosa is a constant challenge. This process is governed largely by local compartmentalised macrophage and dendritic cell (DC) populations which sample incoming environmental antigens prior to trafficking to airways draining lymph nodes for the generation of immune responses (21). Consequently, the respiratory tract requires tightly regulated and efficient mechanisms to facilitate rapid replenishment of these populations following perturbation. Previous studies have identified a reservoir of tissue-resident hematopoietic progenitors localised within the extravascular lung of adult rodents (22, 23), which may serve to fulfil this purpose, whilst simultaneously providing support for the rapid local generation of functionally-specific immune populations during acute inflammatory challenge.

In the present study, we extend current understanding of the developmental kinetics of lung hematopoietic progenitors by comparison to phenotypically and developmentally equivalent HSPC within BM. We demonstrate that HSPC populations are present within the lungs from E18.5 until adulthood, and display microenvironmental-specific population dynamics in regard to subset-specific dominance, compared to their BM counterparts. Moreover, we demonstrate a significant but transient perturbation in this lung HSPC pool following Influenza A viral infection. These findings could have significant implications in how we understand immunological protection within the lungs.

## Results

### Developmental kinetics of LSK subsets within the bone marrow and lung

To map the developmental kinetics of HSPC subsets (Figure S1), paired BM and lung where phenotypically characterised at 6 timepoints spanning E18.5 to adulthood. Primary investigation of the BM identified the classically reported LSK subset (refer to Figure S2 and (24) for gating), which could be clearly partitioned into Flt3^-^CD34^-^ long-term HSC (LT-HSC), Flt3^-^CD34^+^ short-term HSC (ST-HSC) and Flt3^+^CD34^+^ MPP sub-phenotypes (4, 5) (Figure 1A, top panels). Parallel analysis revealed phenotypically equivalent populations of definitive LSK cells residing within the lungs of all age groups examined (Figure 1A, bottom panels; refer to Figure S3 and Figure S4 for gating), supporting previous studies in adult mice (22).

**Figure 1.**
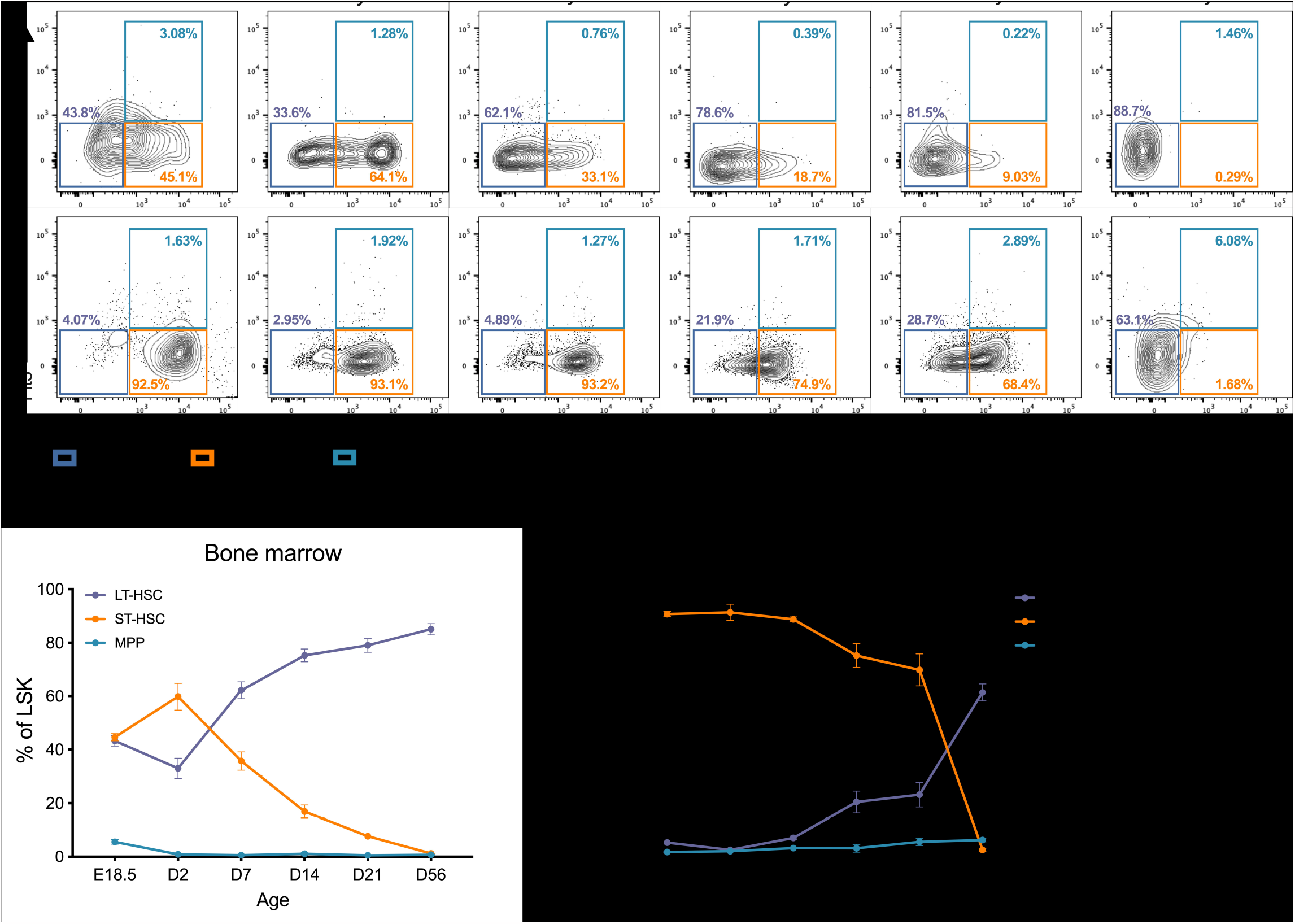
Frequency of LSK subsets within the bone marrow and lung over the course of development. **(A)** Representative flow cytometry plots displaying proportions of long-term HSC (LT-HSC), short-term HSC (ST-HSC) and multipotent progenitors (MPP) within the BM and lung from E18.5 to D56. **(B)** LT-HSC, ST-HSC and MPP kinetics as a proportion of LSK cells within the BM. **(C)** LT-HSC, ST-HSC and MPP as a proportion of LSK cells within the lung. Data are displayed as mean ± SEM.

Assessment of LSK subsets in the BM and lung revealed vastly divergent trajectories. In late gestation (E18.5), the BM LSK compartment was comprised of a relatively equal distribution of LT- and ST-HSC (Figure 1B and Figure S5A). During the first week of life, a progressive shift was observed with the establishment of a LT-HSC dominated BM niche, constituting 62.23% ± 3.20% (mean ± SEM) of the total LSK population by late infancy (D7) and expanding to 85.08% ± 2.11% (mean ± SEM) by adulthood (D56; Figure 1B). The comparatively minor MPP subset remained relatively stable within the BM across all ages assessed (Figure 1B). In contrast to the BM, characterisation of the embryonic (E18.5) lung LSK compartment revealed a ST-HSC dominated niche of 90.0% ± 0.96% (mean ± SEM), which was maintained until weaning at D21 (Figure 1C and Figure S5B). This was proceeded by a dramatic shift in the LSK niche at D56 post-birth, whereby the subpopulation characteristics within both the BM and lung converged in adulthood, with a dominant LT-HSC reservoir at both sites (Figure 1B and C). In parallel with the BM, however, the proportion of lung MPP remained relatively minor and stable throughout development (Figure 1C).

### Developmental kinetics of myeloid progenitors within the bone marrow and lung

Having confirmed the presence of a definitive LSK reservoir within murine lungs throughout life, we next sought to characterise the extravascular myeloid progenitor (MP) compartment. All MP subsets were clearly identifiable within paired BM and lung samples from E18.5 to adulthood (Figure 2A; refer to Figures S2-S4 for gating).

**Figure 2.**
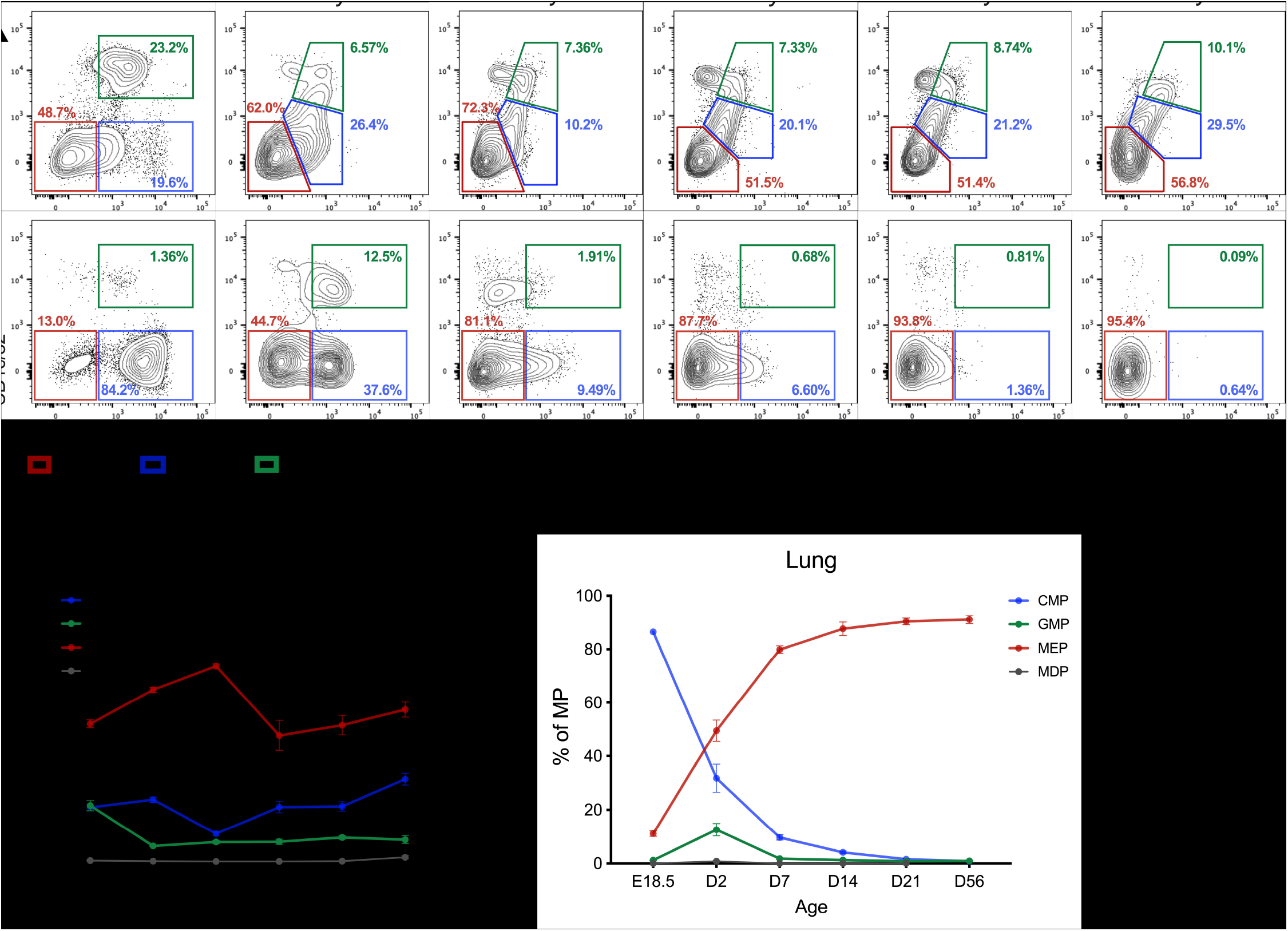
Developmental kinetics of MP subsets within the bone marrow and lung over the course of development. **(A)** Representative flow cytometry plots displaying proportions of common myeloid progenitors (CMP), granulocyte-macrophage progenitors (GMP) and megakaryocyte-erythrocyte progenitors (MEP) within the BM and lung from E18.5 to D56. **(B)** CMP, GMP, MEP and macrophage-dendritic cell progenitor (MDP) kinetics as a proportion of myeloid progenitors (MP) within the BM. **(C)** CMP, GMP, MEP and MDP as a proportion of MP within the lung. Data are displayed as mean ± SEM.

Within the BM, a consistent increase in MEPs was observed during the first week of life, followed by a return to E18.5 levels at D14 and establishment of a MEP pool comprising approximately 50% of BM MPs into adulthood (Figure 2B). In contrast, a transient reduction in CMPs was observed from D2 to D7 (Figure 2B), proceeded by a gradual increase, subsequently exceeding that of E18.5, from D14 until adulthood. BM GMPs demonstrated an initial reduction within the first 48 hours of life compared to embryonic levels, which stabilised at 6-10% of BM MPs throughout life (Figure 2B). While MDPs were able to be identified within the BM of all age groups, they constitute a minor subset of less than 0.5% of MPs across all ages examined (Figure 2B), as previously recognised (25). Further analysis revealed significantly greater Flt3 receptor expression on adult MDPs compared to all other ages (Figure S5C), highlighting the increased requirement for Flt3 ligand (and consequent negative feedback on Flt3 expression) during early postnatal establishment of peripheral DC populations (25). Owing to the relatively stable proportions of MP subsets within the BM over time (Figure 2B), a gradual increase in the totality of all subsets was observed during development, reflective of the increase in total BM cellularity with increasing age (Figure S5D and E). This uniformity was clear upon unbiased clustering (UMAP) of lineage negative BM cells when accounting for age (Figure S5F). We were able to further classify CLPs and pre-NKp, with rapid expansion of BM pre-NKp observed during the first 48 hours of life and peaking at D7, at which point maintenance of a relatively constant population was achieved (Figure S5G). In contrast, BM numbers of CLPs and lung CLPs and pre-NKp remain comparatively stable throughout normal development (Figure S5G and H).

Within the lung MP pool, we observed a striking transition characterised by a rapid switch from a CMP dominated niche at E18.5, to the establishment of an extravascular MEP reservoir by D7 (Figure 2C). The unique prenatal lung HSPC niche was further exemplified by separate clustering of E18.5 lineage negative cells compared to all other ages examined during unbiased analysis (Figure S5I). A rapid and transient increase in lung GMP was observed within the first 48 hours of birth (Figure 2C). However, this GMP pool had returned to >2% of the MP pool by D7 where it was maintained throughout life (Figure 2C). In parallel with the BM, MDP within the lung constituted a relatively minor population (Figure 2C). Due to the low numbers in adult lung, we were unable to evaluate lung MDP Flt3 expression (data not shown). Overall, MP comprise a minor niche within the adult homeostatic lung, and this pattern of immunological development is reflected by a gradual reduction in the total number of CMP, GMP and MDP with increasing age, while MEP remain stable given their requirement for platelet biogenesis (22) (Figure S5J).

As a supplementary analysis to examine whether the phenotypically analogous lung HSPC pool exhibited an equivalent developmental trajectory as the BM niche, we generated diffusion maps (26), which can faithfully reconstruct hematopoietic differentiation (27), on representative samples at each timepoint (Figure S6). Focussing on the root population of HSCs (LSK), we observe early separation into the downstream CMP pool within the BM (Figure S6A), however, lung HSCs exhibit a close association with CMPs from late prenatal development till the first days of life, with divergence of these populations more prominent with increasing age (Figure S6B). As expected, spatial clustering of CMPs, GMPs and MEPs is preserved across all ages examined; reinforcing the observation that HSCs and MPs within the lung emulate the developmental dynamics observed within the BM.

### Influenza A viral infection induces tissue microenvironment-specific HSPC responsiveness

We next sought to determine the tissue microenvironment-specific impact of mouse-adapted A/PR8/34 H1N1 Influenza A virus (IAV) infection on lung cellular dynamics. Adult female BALB/c mice were infected with a sub-lethal dose of IAV and cellular responses assessed during the acute, peak and resolution phases of disease (Figure 3A). As previously observed (28), infected mice exhibit clinical disease scores peaking at day-post-infection (dpi) 6-8 (Figure 3B) and significant weight loss up to 22% ± 0.02% (mean ± SEM; Figure 3C). All infected mice recovered to pre-infection clinical disease scores by dpi 22, however fluctuations in weight gain remained present until at least dpi 32 (Figure 3B and C).

**Figure 3.**
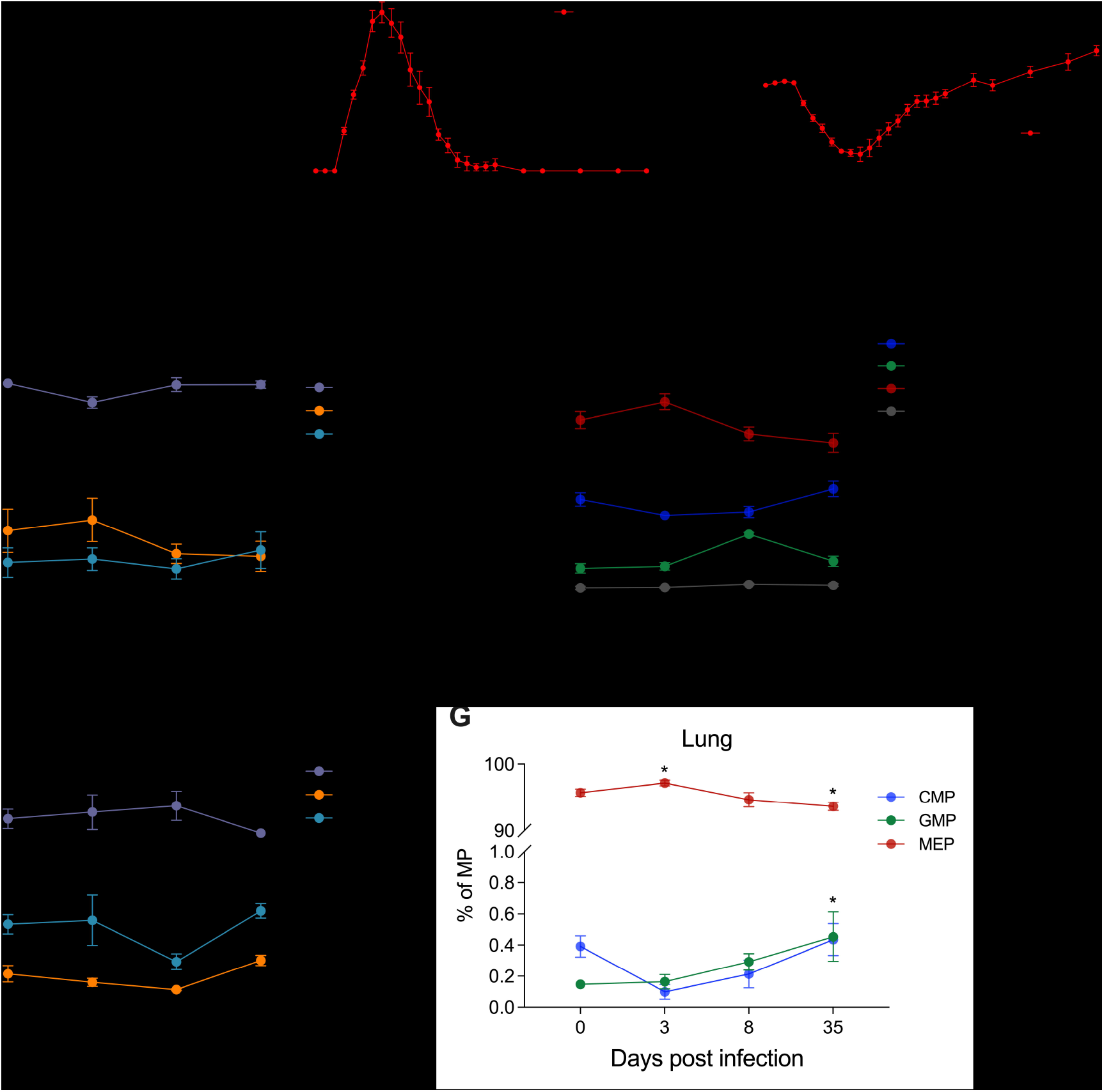
Modulation of HSPC populations in the bone marrow and lung following Influenza A viral infection. **(A)** Female BALB/c mice were infected with mouse-adapted A/PR8/34 H1N1 Influenza A virus (IAV) at 56 days of age (dpi 0) and autopsied at the acute (dpi 3), peak (dpi 8) and resolution (dpi 35) phases of infection. **(B)** Clinical disease scores and **(C)** weight of naïve non-infected (dpi 0) and IAV infected mice. **(D)** LT-HSC, ST-HSC and MPP kinetics as a proportion of LSK cells within the BM of naïve (dpi 0) and IAV infected mice (dpi 3-35). **(E)** CMP, GMP, MEP and macrophage-dendritic cell progenitor (MDP) kinetics as a proportion of myeloid progenitors (MP) within the BM of naïve (dpi 0) and IAV infected mice (dpi 3-35). **(F)** LT-HSC, ST-HSC and MPP kinetics as a proportion of LSK cells within the lungs of naïve (dpi 0) and IAV infected mice (dpi 3-35). **(G)** CMP, GMP, MEP and macrophage-dendritic cell progenitor (MDP) kinetics as a proportion of myeloid progenitors (MP) within the lungs of naïve (dpi 0) and IAV infected mice (dpi 3-35). Data are mean ± SEM of *n*=5 mice per group. Statistical significance was determined using Ordinary Two-way ANOVA with Sidak multiple comparisons test (B and C) or RM Two-way ANOVA with Geisser-Greenhouse correction with Fisher’s LSD test (D-G) and displayed as \**p* < 0.05, \*\**p* < 0.01 and \*\*\**p* < 0.001 compared to dpi 0.

HSPC populations were initially characterised by Uniform Manifold Approximation and Projection (UMAP) to gain a holistic view of the infection dynamics. While global changes in HSPC were challenging to visualise (Figure S7A), infection significantly reduced the proportion of LT-HSC within the BM at dpi 3 when compared to uninfected dpi 0 controls (Figure 3D). Downstream, infection increased BM GMPs at the peak of disease, with a concomitant decrease in CMPs, with both returning to baseline proportions upon disease resolution (Figure 3E). In parallel, infection amplified the CLP (Figure S7B) and pre-NKp (Figure S7C) pools at the acute and peak phases of infection within the BM, with pre-NKp proportions positively correlating with the mature natural killer (NK) cell response at dpi 8 (Figure S7D and E). In the lung, infection resulted in a transient reduction in MPP (Figure 3F) at the peak of disease, while a significant increase in MEP occurred during the acute phase at dpi 3, which subsided upon disease resolution (Figure 3G). Moreover, distinct from the BM response, a continual increase in GMP was detected within the lungs of infected mice, reaching significance at dpi 35 (Figure 3G).

### Influenza A viral infection promotes concomitant pre-pDC and mature pDC infiltration into the lungs at the peak of infection

Finally, unbiased analysis of committed populations highlighted global changes in monocytes, inflammatory dendritic cells (infDC), plasmacytoid DCs (pDC) and NK cells within the lungs (Figure 4A) and to a lesser extent infDCs and pDCs within the BM (Figure S7F) following infection. Within the lungs, infection drove a significant increase in pDC precursors (pre-pDC) with parallel expansion of mature pDCs (Figure 4B and C). A reduction in conventional DC precursors (pre-cDC) was additionally observed at dpi 3 (Figure 4D), potentially reflecting rapid cDC commitment-induced depletion as a mechanism to replenish mature cDCs trafficking to lung draining lymph nodes following acquisition of viral antigen (29), although only a trend in mature cDC reduction was observed during the acute phase (Figure 4E). Furthermore, we identified an early phase rapid increase in infDCs (Figure 4F), known to promote the recruitment of NK cells (Figure S7G) (30) during acute infection. Infected mice additionally experienced an acute reduction in interstitial macrophages, as previously observed (28), prior to repopulation at disease resolution (Figure 3G), potentially via proliferation of lung-resident precursors (31).

**Figure 4.**
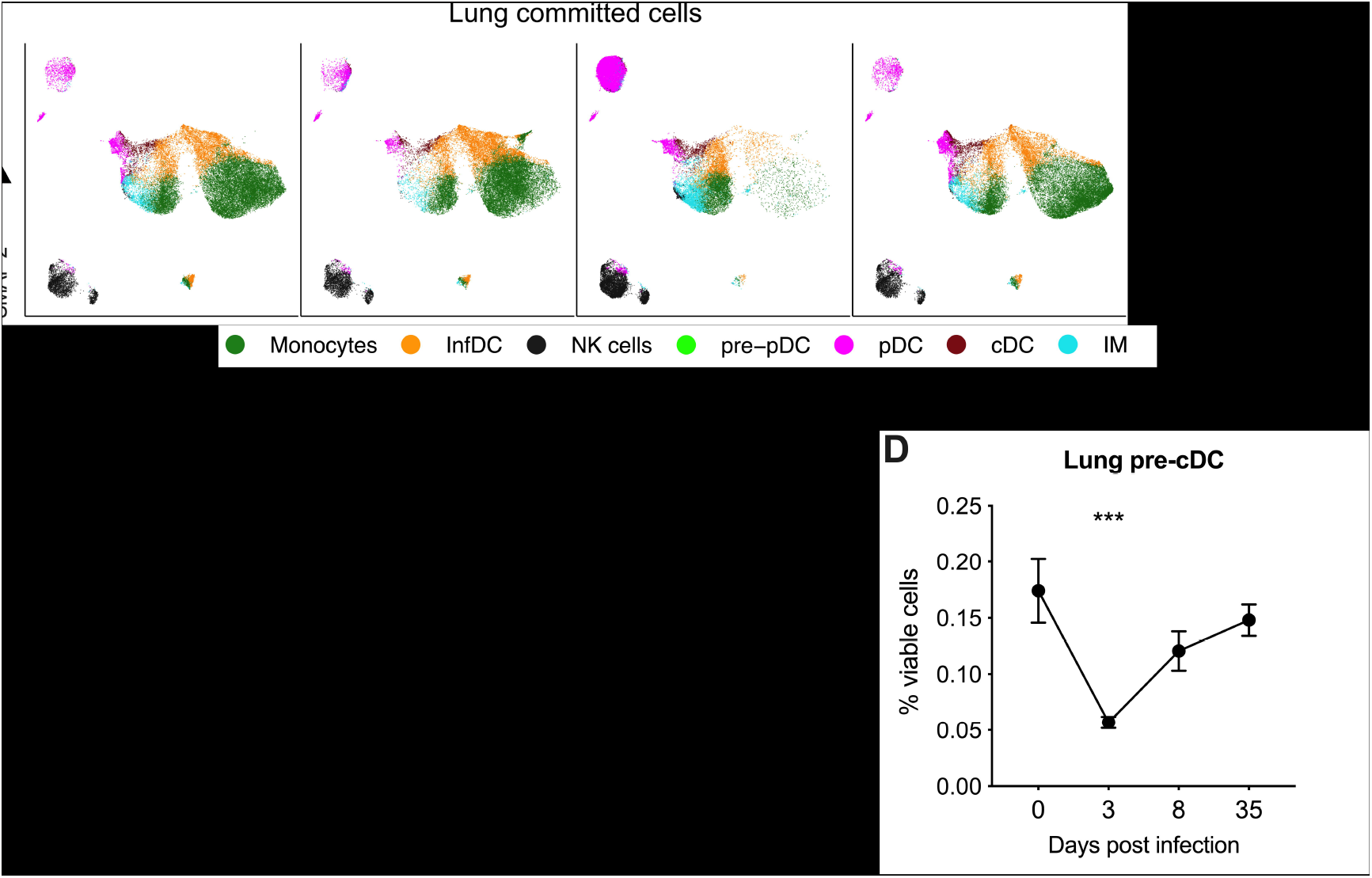
Influenza A viral infection upregulates proinflammatory myeloid responsiveness in the lung. **(A)** UMAP of committed subsets identified in the lungs of naïve and IAV infected mice. Proportions of **(B)** pre-pDC, **(C)** pDC, **(D)** pre-cDC, **(E)** cDC, **(F)** infDC and **(G)** interstitial macrophages within the lungs of naive and IAV infected mice. Data are mean ± SEM of *n*=6 mice per group. Statistical significance was determined using Ordinary one-way ANOVA with Fisher’s LSD test and displayed as \*\**p* < 0.01, \*\*\**p* < 0.001 and \*\*\*\**p* < 0.0001 compared to dpi 0.

## Discussion

Following pioneering studies identifying cells with HSPC characteristics within the lung (22, 23), we have phenotypically mapped HSPC subset ontogeny in the lungs and BM of mice from embryonic day 18.5 to adulthood, to gain further insight into their developmental kinetics. Consistent with previous reports (22), we demonstrate the existence of a definitive extravascular pool of lung HSPC and associated myeloid and lymphoid progenitors phenotypically equivalent to their BM counterparts, which display unique age-dependent and microenvironment-specific population dynamics. Moreover, we reveal global preservation of the classic hierarchical structure of HSPC by mapping lineage commitment trajectories, validating the developmental kinetics of this unique extramedullary niche.

During prenatal development, the fetal lung experiences a dramatic influx of liver-derived monocytes capable of rapidly establishing a resident colony of alveolar macrophages by the first week of life (18, 32). Our current findings demonstrating the presence of all major MP subsets in the lungs at E18.5, with an abundance of CMPs, suggests these prenatal seeding events extend to include more definitive populations. A limitation of this study is that as vascularisation of fetal long bones is initiated at E16 (33), and perfusion of lung vasculature during the saccular stage is not possible due to tissue fragility, we have potentially not explicitly excluded cells in circulation at E18.5 and D2. Nevertheless, the transient spike in postnatal GMP on D2 closely resembles the population dynamics of lung granulocytes, known to temporarily peak during the first days of life (34). Early life is a highly susceptible period owing to developmental-associated deficiencies in immune regulatory functions that mature postnatally, exemplified by the inherently delayed neonatal neutrophilic response due to an insufficient BM pool, impaired chemotactic activity and reduced capacity for tissue extravasation (35). As such, a transient spike in local progenitors capable of rapid neutrophil generation may signify an intrinsic innate mechanism designed to facilitate early postnatal protection against bacterial infection within the lungs. Moreover, immune maturation is directly related to gestation age at birth (36), and perturbation of this early postnatal GMP pool as a result of prematurity may subsequently contribute towards the heightened state of vulnerability in these infants (37).

An array of specialised local precursors capable of multilineage differentiation are known to populate the adult lung and airways (38, 39). However, a *bona fide* niche of adult lung HSPCs and MPs has only recently been demonstrated (22). Together with lung resident mesenchymal stromal cells (40), the presence of which is a fundamental constituent ensuring HSPC survival and proliferation (41), provides credence to a microenvironment supportive of a peripheral HSPC niche. Our current findings compliment the existence of a specialised adult lung haematopoietic progenitor niche, mirroring the MEP-dominated population dynamics originally described (22). It is pertinent to note that lung CMPs, GMPs and MEPs in the current study are phenotypically analogous to those found in fetal BM, suggesting these cells exhibit an immature phenotype, a feature previously recognised with regards to megakaryocyte profiles in the adult lung (22). This hematopoietic progenitor niche may therefore represent a quiescent pool under homeostatic conditions, given the established roles for bipotential alveolar progenitors and club cells in the maintenance of airways homeostasis (14, 15). However, with 70m^2^ of lung in constant exposure to environmental insults, we speculate this progenitor pool provides a central function in repopulating the lungs both during and following infectious perturbation or injury (23).

In this regard, our observation of increased lung MEPs during the acute phase of IAV infection potentially highlights a state of reinforced megakaryopoiesis required for platelet generation as a result of viral-indued thrombocytopenia, as recently observed within the BM following IAV infection (42). Furthermore, we demonstrate a reduction in adult lung MPPs following acute Influenza A viral infection, with concomitant upregulation of local pDC, NK cell and infDC effector populations during the acute and peak phases of infection, suggesting lung progenitors may in fact contribute towards local inflammatory responses prior to ensuing support from the BM. Indeed, BM hematopoietic activity preferentially shifts towards granulopoiesis in response to infection to compensate for enhanced mobilisation and trafficking towards the primary site of challenge (43). This was observed within the BM GMP pool in this study following infection, and previously in response to allergic airways challenge (44) and in patients with severe acute respiratory syndrome-coronavirus-2 (SARS-CoV-2) infection (45). It is important to note we are currently unable to determine whether the transient reduction in lung MPPs is a direct result of active commitment, or whether such progenitors are susceptible to IAV infection, as observed with alveolar progenitors (46). Nevertheless, repopulation of lung progenitor subsets is evident upon disease resolution, with an overcompensation of GMPs, illustrating the inherent capacity of this specialised progenitor niche to re-establish homeostatic cellular population dynamics following significant local perturbation. However, given the continuous exposure to environmental stress, and heightened susceptibility of MPs to stress-induced exhaustion (47), this unique lung HSPC pool may experience limited self-renewal capacity with increasing age.

In conclusion, the results of this study complement the paradigm shift in how we understand the origins and maintenance of local immune control and defence mechanisms in the lungs, with growing recognition that the lungs have significant hematopoietic and self-renewal potential. A better understanding of the role of these cells will have significant implications for how we approach prevention of inflammatory lung diseases, and promote the return to normal homeostasis after lung injury.

## Acknowledgements

The authors wish to acknowledge the animal technicians at the Telethon Kids Institute Bioresources Centre.

## Author Contributions

K.T.M, J.-F.L.-J and D.H.S designed the study. K.T.M, J.-F.L.-J and J.F.R performed the experiments and analysed the data. P.G.H and P.A.S contributed to discussions on data interpretation. K.T.M, P.A.S and D.H.S wrote the manuscript. All authors reviewed the final version of the manuscript.

## Methods

### Animals

Specified pathogen-free BALB/c mice were purchased from the Animal Resource Centre (Murdoch, Western Australia). All mice were housed under specific pathogen-free conditions at the Telethon Kids Institute Bioresources Facility under a 12-hour light/12-hour dark cycle with access to food and water *ad libitum*. In-house-bred BALB/c offspring of both sexes were used in all studies, with the exception of IAV infected mice, which were all females.

### Time-mated pregnancies

Female BALB/c mice 8-12 weeks of age were time-mated with male BALB/c studs 8-26 weeks of age. Male studs were housed separately in individual cages with 1-2 females overnight. The presence of a vaginal plug the following morning was used as an indicator of mating. The day of vaginal plug detection was designated embryonic (E) day 0.5.

### Influenza A H1N1 viral infection

Mouse-adapted Influenza A/Puerto Rico/8/1934 (PR8) virus (IAV) from the American Type Tissue Culture Collection was prepared in allantoic fluid of 9-day-old embryonated hen eggs. Stock virus was sub-passaged through Madin-Darby canine kidney (MDCK) cells in Dulbecco’s modified Eagle’s medium (DMEM; Gibco), harvested as tissue culture supernatant and viral titres determined by cytopathic effects on MDCK cells and expressed as the mean log_10_ tissue culture infective dose required to kill 50% of the cells (TCID_50_) over a 5-day incubation period (28). Female BALB/c mice at 56 days of age were intranasally inoculated with 20 TCID_50_ PR8 in 25μl phosphate-buffered saline (PBS) as previously described (48). IAV infected mice were monitored daily for weight loss and clinical score from day-post-infection (dpi) 0 – 19, and then twice-weekly until the end of experiment.

### Tissue collection

#### Fetuses

Fetal tissue was collected as previously described (24). Briefly, pregnant BALB/c mice were sacrificed on E18.5 and both horns of the uterus were removed before fetuses (*n* = 17) were humanely sacrificed by decapitation. Fetal hind legs and whole lungs were removed and collected into cold PBS + 0.1% bovine serum albumin (BSA) and stored on ice. Dead fetuses were excluded from the study. *D2 mice:* Mice were sacrificed at 2 days of age (*n* = 8). Femur and tibia (long bones) from both hind legs and whole lungs were removed and collected into cold PBS + 0.1% BSA and stored on ice. *D7, D14, D21 and D56 mice:* Mice were sacrificed at 7 (*n* = 9), 14 (*n* = 6), 21 (*n* = 5) and 56 (*n* = 12) days of age. Lungs were perfused via cardiac flush with cold PBS + 0.1% BSA. Lungs and long bones were collected in cold PBS + 0.1% BSA and stored on ice. All lung tissue was weighted at the time of collection. *IAV infected mice:* Infected mice were sacrificed at dpi 0 (*n* = 5), 3 (*n* = 5), 8 (*n* = 5) and 35 (*n* = 5). Lungs and long bones were collected as above.

### Single-cell suspension preparation

#### Lungs and E18.5/D2 BM

Lung and BM samples were prepared by mincing with a scalpel followed by enzymatic digestion as previously described (44). Briefly, minced tissues were resuspended in GKN + 10% fetal calf serum (FCS) with collagenase IV (Worthington Biochemical Corporation) and DNase (Sigma-Aldrich) at 37°C under gentle agitation for 60 minutes (BM) or 90 minutes (lung). Digested samples were filtered through sterile nylon. Cell suspensions were centrifuged, and pellets resuspended in red blood cell (RBC) lysis buffer. Cells were washed with PBS and pelleted. Supernatant was removed, and pellet resuspended in PBS + 0.1% BSA for total cell counts. *D7, D14, D21, D56 and IAV infected mice BM:* Long bones were flushed with GKN + 5% FCS using a 25-guage needle. Cells were disaggregated and filtered through sterile nylon. Cell suspensions were centrifuged, and pellets resuspended in RBC lysis buffer. Cells were washed with PBS and pelleted. Supernatant was removed, and pellet resuspended in PBS + 0.1% BSA for total cell counts.

### Flow cytometry

Single-cell suspensions were used for all immunostaining. Panels of monoclonal antibodies were developed to enable immunophenotypic characterisation of hematopoietic stem and progenitor cells (Table S1) and committed cells (Table S2). Data acquisition was performed on a 4-laser LSR Fortessa (BD Bioscience). All samples were kept as individuals and not pooled. Manual immunophenotypic characterisation was performed using FlowJo software (version 10.7.1, BD Biosciences) and HSPC gating strategies outlined as previously described (24) (fetal BM analysis) and in Figures S2-S4. Committed cell gating strategy used was as previously described (44). Fluorescence minus one staining controls were used where required.

### Unsupervised flow cytometry analysis

Unsupervised analysis was performed in the R environment for statistical computing (version 3.6.2). All data underwent initial pre-processing to remove cellular debris, doublets and non-viable cells. Cleaned data was manually gated on Lin^−^ (HSPC) or CD3^−^ CD19^−^ CD11b^+/ −^ B220^+/ −^ (committed) cells, prior to down-sampling to the maximum number of events present within the sample with the least number of events (Lung HSPC = 13,494 events; Lung committed = 7,562 events; BM HSPC = 9,870 events; BM committed = 62,950 events). Data underwent logicle transformation followed by dimensionality reduction using the *CATALYST* package (49), employing initial high-resolution *FlowSOM* clustering prior to low-resolution metacluster generation via *ConsensusClusterPlus*. Uniform Manifold Approximation and Projection (UMAP) (50) was performed on 500 cells per sample. Heatmaps displaying antigen expression within each defined UMAP cluster were used to immunophenotypically characterise individual clusters.

### Diffusion map analysis

Diffusion mapping was performed in the R environment for statistical computing (version 3.6.2) with the *destiny* package available via Bioconductor (51). Three representative samples were selected for each tissue and timepoint. Samples underwent pre-processing (compensation, gating) and logicle transformation. Samples were enriched for HSCs, (LT-HSC and ST-HSC combined), CMPs, GMPs, and MEPs, as determined by manual characterisation with FlowJo software. Random down-sampling to equal cell numbers across tissue and timepoints was applied where possible, to limit the influence of cellular composition. This resulted in 150 cells per cell type per tissue/timepoint available for analysis, with two exceptions; 1) D21 lung samples were down-sampled to 59 cells to accommodate lower cell numbers for GMPs (n=59), CLPs (n=77), and MEPs (n=129), 2) D56 lung samples were largely absent of GMP (n=4) and CMP (n=6) cells. Diffusion maps were defined by the expression of CX3CR1, c-Kit, Sca-1, NKG2D, CD34, Flt3, IL-7Rα, and CD16/32.

### Statistical analysis

Statistical analyses and graphing were performed using GraphPad Prism (version 8.3.0, GraphPad Software). Statistical significance of *p > 0*.*05* was considered significant. Ordinary one-way analysis of variance (ANOVA) with Uncorrected Fisher’s LSD test was used for immunophenotypic analysis. Clinical response significance was determined using two-way ANOVA with Sidak multiple comparisons test. Linear correlation testing was determined using Pearson correlation (r) test.

### Study approval

All animal experiments were formally approved by the Telethon Kids Institute Animal Ethics Committee, which operates under guidelines developed by the National Health and Medical Research Council of Australia for the case and use of animals in scientific research.

