## Supplementary data for "Mapping lung hematopoietic progenitors: Developmental kinetics and response to Influenza A viral infection"

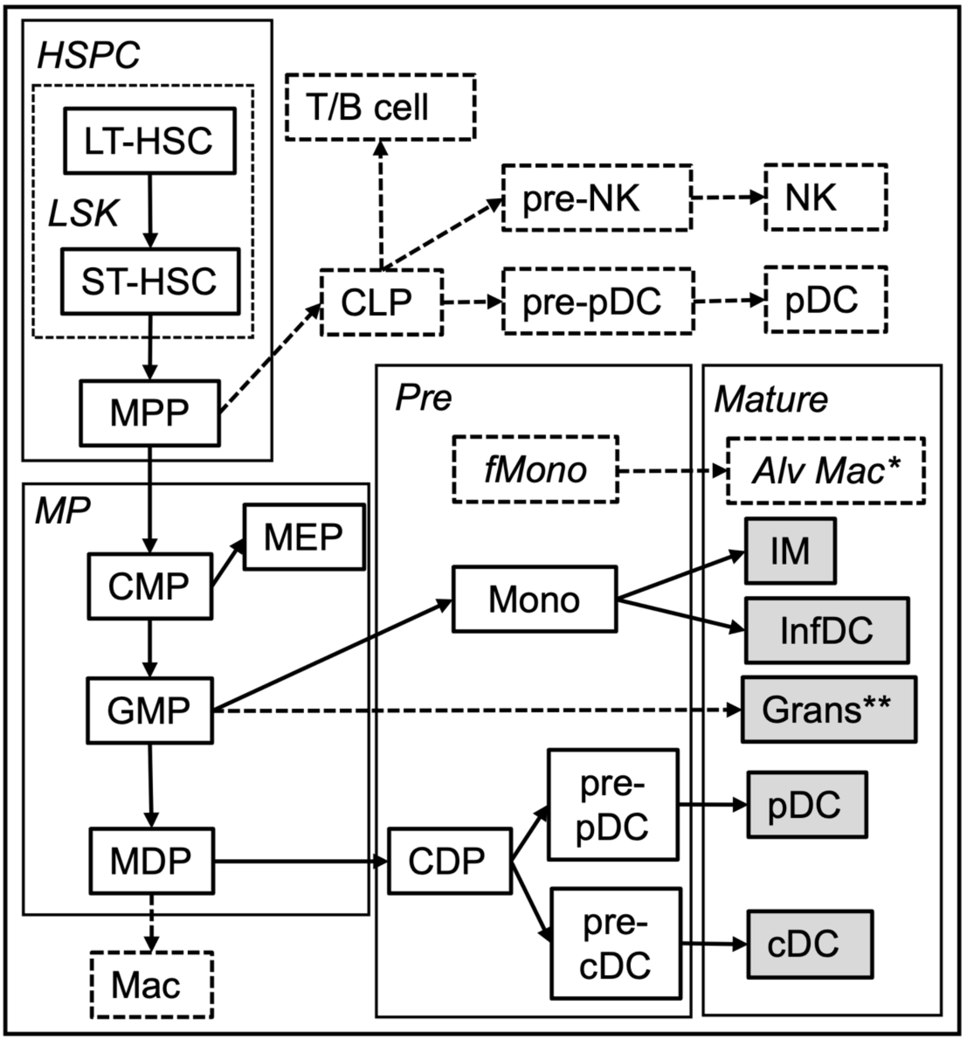


**Figure S1.** Myeloid progenitor cell development in bone marrow and lung tissue. ***HSPC*:** Haematopoietic Pluripotent Stem Cell; ***LSK*:** $Lin^{-}Sca\text{-}1^{+}c\text{-}Kit^{+}$; **LT-HSC:** Long-term Haematopoietic Stem Cell; **ST-HSC:** Short-term Haematopoietic Stem Cell; **MPP:** Multipotent progenitor; **CLP:** Common Lymphoid Progenitor; ***MP***: Myeloid Progenitors; **CMP:** Common Myeloid Progenitor; **MEP:** Megakaryocyte-Erythrocyte Progenitor; **GMP:** Granulocyte-Monocyte Progenitor; **Grans:** Granulocytes; **MDP:** Macrophage-Dendritic Cell Progenitor; **CDP:** Common Dendritic Cell Precursor; ***Pre:*** Lung tissue precursors; **fMono:** Fetal Monocyte; **Mono:** Monocyte; **pre-pDC:** pre-plasmacytoid dendritic cell; **pre-cDC:** pre-Conventional Dendritic Cell; ***Mature:*** Mature lung tissue myeloid cells; **Alv Mac**: Alveolar Macrophage; **IM:** Interstitial Macrophages; **InfDC**: Inflammatory DC **pDC:** plasmacytoid Dendritic Cell; **cDC:** conventional Dendritic Cell. **Mouse alveolar macrophages are derived from a separate lineage of fetal-origin monocytes but may be replaced by recruitment of BM-derived monocytes into the alveolar niche following lung injury. ** Includes neutrophils, eosinophils, and basophils via myeloblast/myelocyte progenitors*

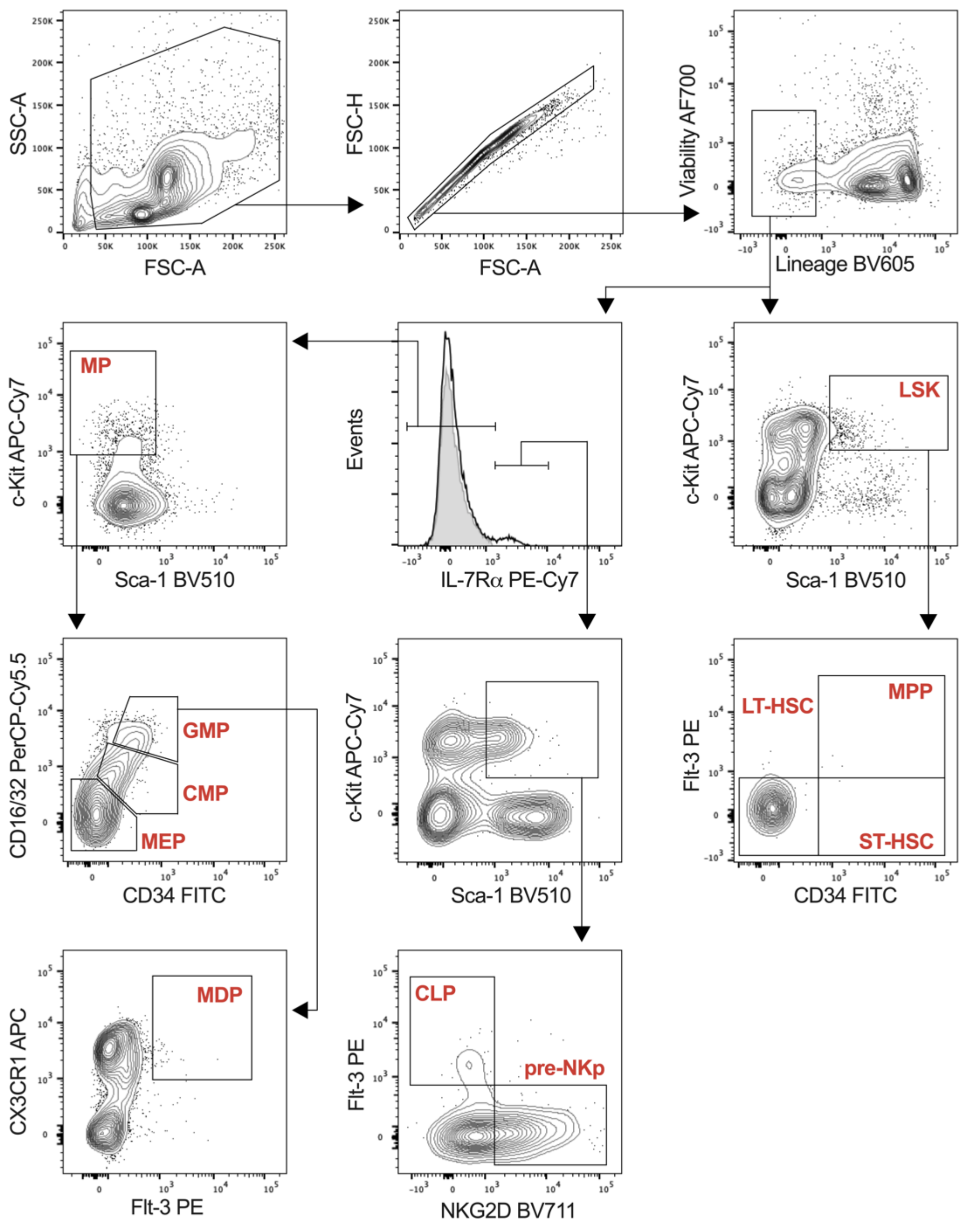


**Figure S2. Postnatal bone marrow HSPC gating strategy.** Gating strategy used to immunophenotypically characterise HSPC populations within postnatal BALB/c bone marrow. Shaded area of the histogram represents IL-7Rα FMO control.

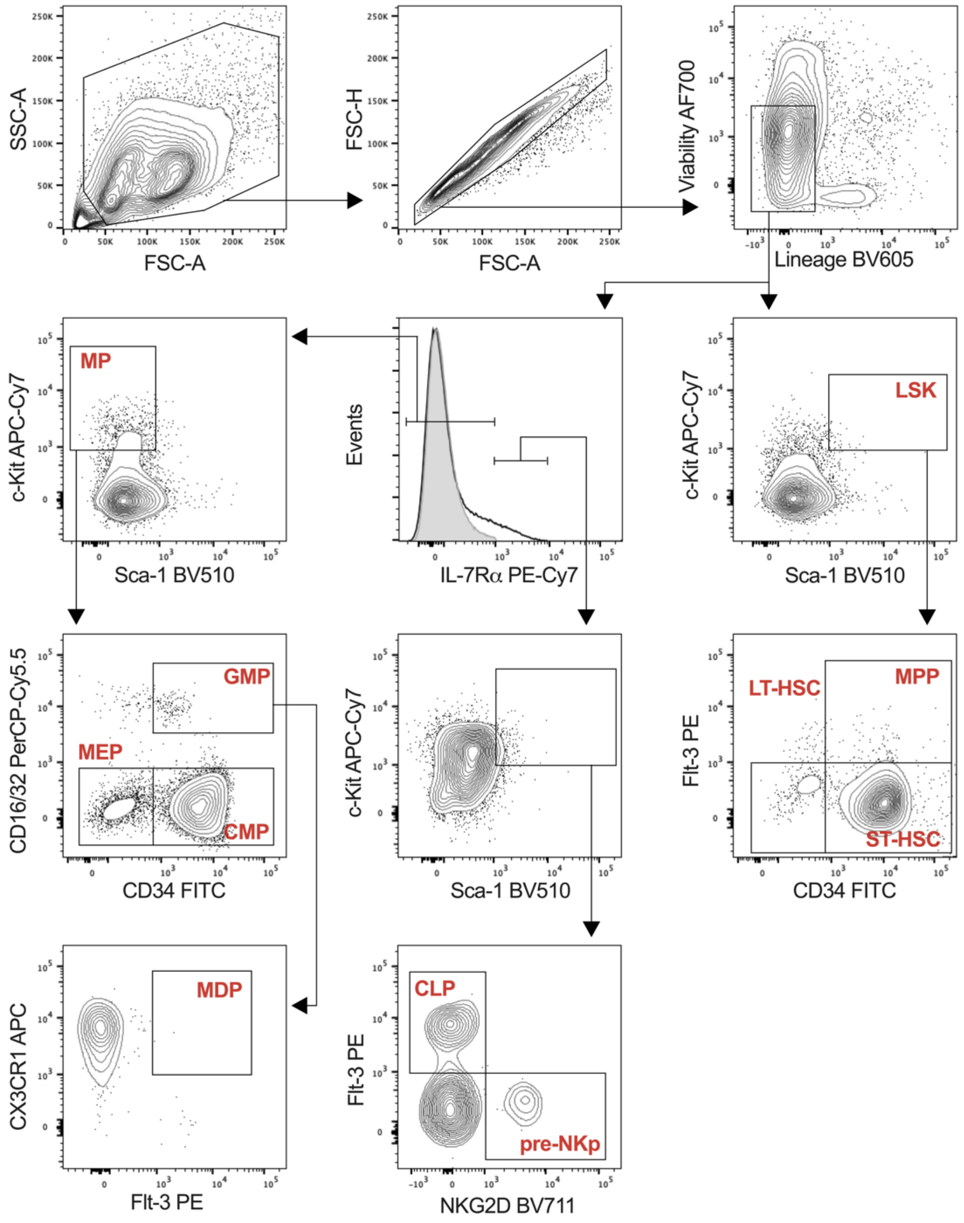


**Figure S3. Fetal lung HSPC gating strategy.** Gating strategy used to immunophenotypically characterize HSPC populations within fetal lungs. Shaded area of the histogram represents IL-7Rα FMO control.

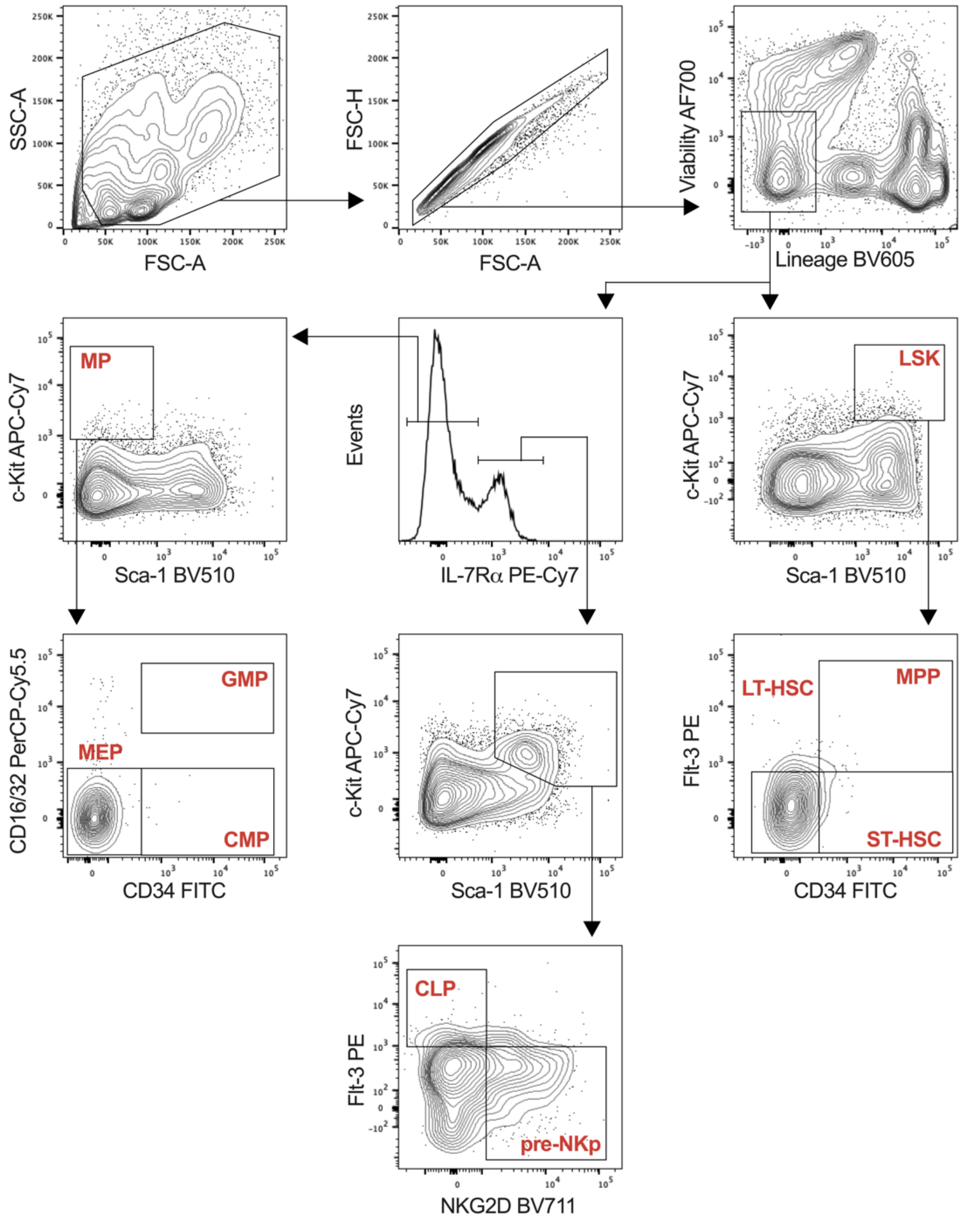


**Figure S4. Postnatal lung HSPC gating strategy.** Gating strategy used to immunophenotypically characterize HSPC populations within postnatal BALB/c lungs.

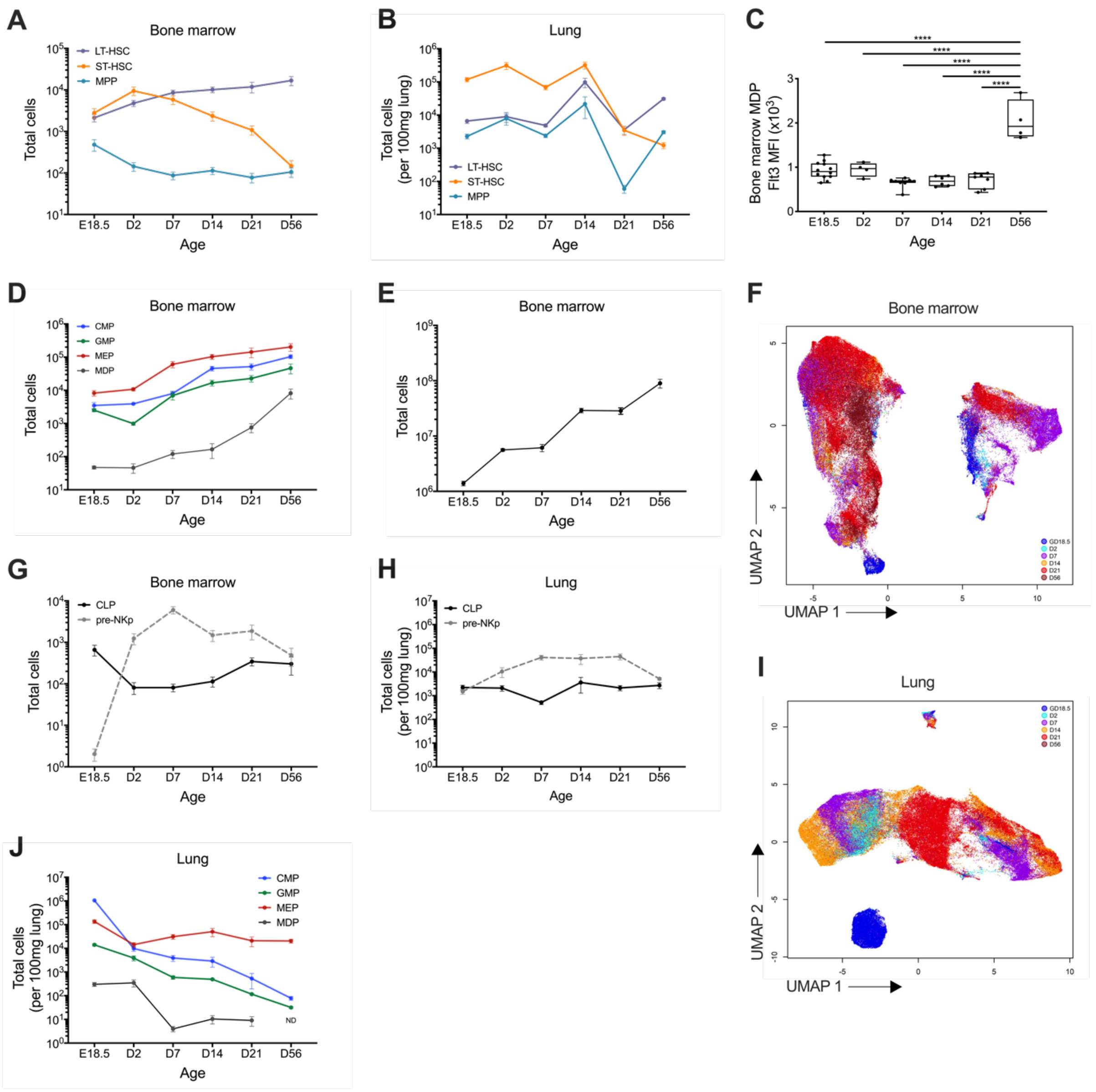


**Figure S5. Total HSPC subsets within bone marrow and lungs. (A)** Absolute number of LT-HSC, ST-HSC and MPP within the BM. **(B)** Total number of LT-HSC, ST-HSC and MPP per within 100mg of lung tissue. **(C)** Mean fluorescence intensity (MFI) of Flt3 expression on MDP within the BM from E18.5 until 56 days of age. **(D)** Absolute number of CMP, GMP, MEP and MDP within the BM. **(E)** Total cellularity of the BM from E18.5 until 56 days of age. **(F)** UMAP depicting total Lin$^-$ cells within the bone marrow for all age groups sampled. **(G)** Absolute number of CLP and pre-NKp within the BM. **(H)** Total number of CLP and pre-NKp within 100mg lung tissue. **(I)** UMAP depicting total Lin$^-$ cells within the lung for all age groups sampled. **(J)** Total number of CMP, GMP, MEP and MDP per 100mg of lung tissue. Data are from individual animals and displayed as line plot showing mean ± SEM (A-B, D-H) or box and whisker plots showing minimum to maximum values (C). Statistical significance was determined using Ordinary one-way ANOVA with Fisher's LSD test and displayed as ****$p < 0.0001$.

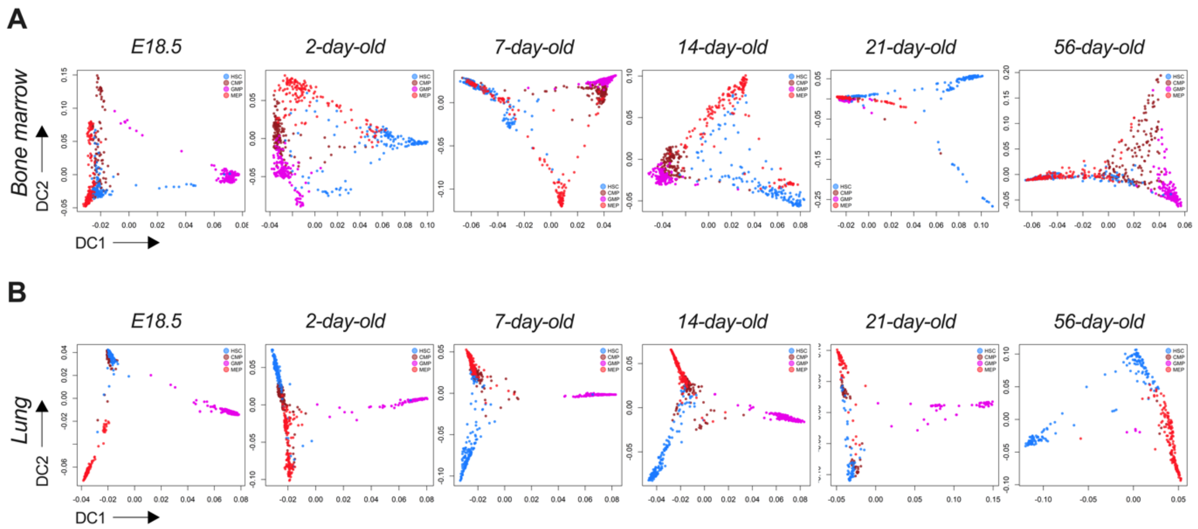


**Figure S6. Extravascular lung myeloid progenitors parallel developmental kinetics observed in the bone marrow.** Diffusion maps of selected cell types from the **(A)** BM and **(B)** lung from E18.5 to D56. Samples were down-sampled to the least abundant terminal cell type prior to analysis, where possible.

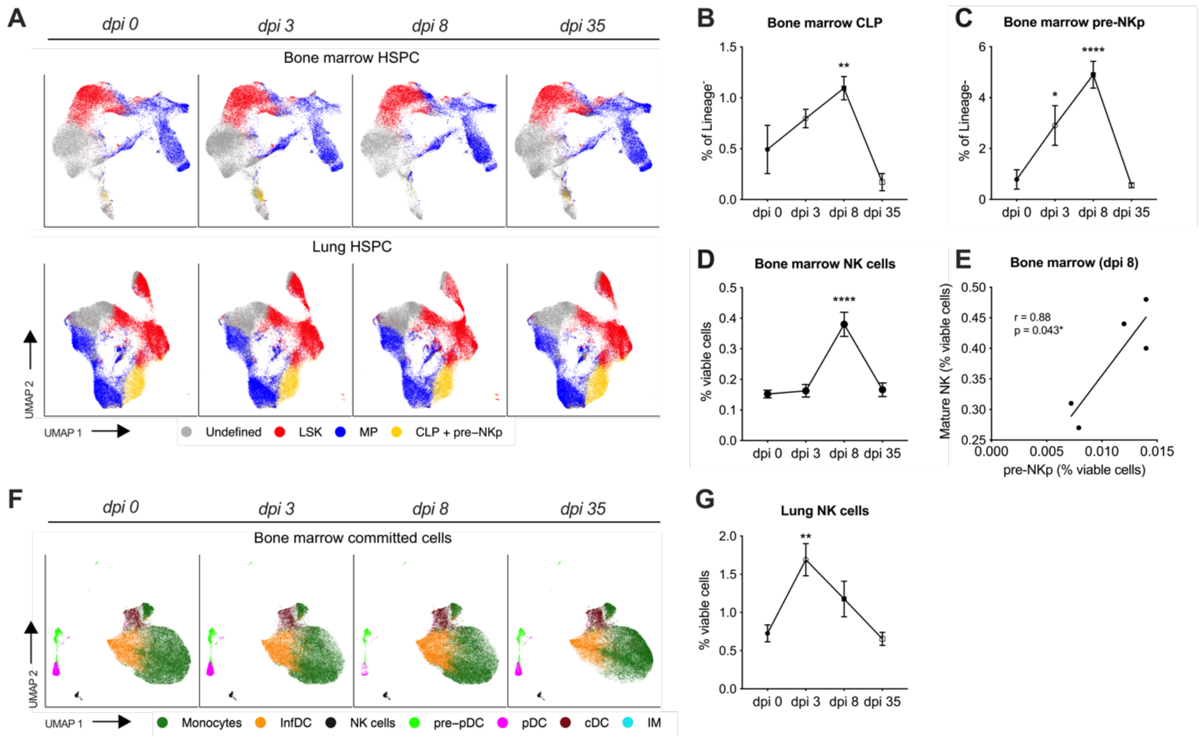


**Figure S7. Impact of IAV infection on HSPC and committed cell populations in the bone marrow and lungs. (A)** UMAP of $Lin^-$ cells displaying LSK, myeloid progenitor (MP), common lymphoid progenitor (CLP) and pre-natural killer cell progenitor (pre-NKp) and undefined clusters in BM and lung naïve (dpi 0) and IAV infected mice (dpi 3-35)**.** Proportions of **(B)** CLP, **(C)** pre-NKp and **(D)** NK cells within the BM of naïve and IAV infected mice. **(E)** Linear correlation between the proportion of NK cells and pre-NKp at dpi 8 within BM of IAV infected mice. **(F)** UMAP of committed subsets identified in the BM of naïve and IAV infected mice. **(G)** Proportion of NK cells within the lungs of naive and IAV infected mice. Data are mean ± SEM of *n*=6 mice per group. Statistical significance was determined using Ordinary one-way ANOVA with Fisher's LSD test (B-D and G) and displayed as $^{**}p < 0.01$ and $^{****}p < 0.0001$ compared to dpi 0. Correlations (E) were determined by Pearson test.

**Table S1. Hematopoietic stem and progenitor cell flow cytometry panel**

| Marker | Fluorochrome | Clone | Supplier |
|---|---|---|---|
| CD2 | Biotin | RM2-5 | BD Biosciences |
| CD3 | Biotin | 145-2C11 | BD Biosciences |
| CD4 | Biotin | GK1.5 | BD Biosciences |
| CD5 | Biotin | 53-7.3 | BD Biosciences |
| CD8a | Biotin | 53-6.7 | BD Biosciences |
| CD19 | Biotin | 1D3 | BD Biosciences |
| B220 (CD45R) | Biotin | RA3-6B2 | BD Biosciences |
| Gr-1 | Biotin | RB6-8C5 | BD Biosciences |
| Ter119 | Biotin | TER-119 | BD Biosciences |
| CD16/32 | PerCP-Cy5.5 | 2.4G2 | BD Biosciences |
| CD34 | FITC | RAM34 | BD Biosciences |
| IL-7Ra (CD127) | PE-Cy7 | SB/199 | BD Biosciences |
| Flt3 (CD135) | PE | A2F10.1 | BD Biosciences |
| c-Kit (CD117) | APC-Cy7 | 2B8 | BD Biosciences |
| Sca-1 | BV510 | D7 | BD Biosciences |
| CX3CR1 | APC | SA011F11 | BioLegend |
| NKG2D | BV711 | CX5 | BD Biosciences |
| Viability | Alexa Fluor 700 | - | BD Biosciences |
| Streptavidin | BV605 | - | BD Biosciences |

**Table S2. Committed cell flow cytometry panel**

| Marker | Fluorochrome | Clone | Supplier |
|---|---|---|---|
| CD3 | FITC | 17A2 | BD Biosciences |
| CD11b | BV510 | M1/70 | BD Biosciences |
| CD11c | BV711 | HL3 | BD Biosciences |
| CD19 | APC-H7 | 1D3 | BD Biosciences |
| B220 (CD45R) | PerCP-Cy5.5 | RA3-6B2 | BD Biosciences |
| Gr-1 | Biotin | RB6-8C5 | BD Biosciences |
| NKp46 | PE-Cy7 | 29A1.4 | BioLegend |
| SIRPa (CD172) | APC | P84 | BioLegend |
| I-A/I-E | BV421 | M5/114.15.2 | BioLegend |
| F4/80 | BV785 | BM8 | BioLegend |
| Viability | Alexa Fluor 700 | - | BD Biosciences |
| Streptavidin | BV605 | - | BD Biosciences |
